# Transition metal binding selectivity in proteins and its correlation with the phylogenomic classification of the cation diffusion facilitator protein family

**DOI:** 10.1101/164509

**Authors:** Shiran Barber-Zucker, Boaz Shaanan, Raz Zarivach

## Abstract

Divalent d-block metal cations (DDMCs), such as Fe, Zn and Mn, participate in many biological processes. Understanding how specific DDMCs are transported to and within the cell and what controls their binding selectivity to different proteins is crucial for defining the mechanisms of metalloproteins. To better understand such processes, we scanned the RCSB Protein Data Bank, performed a *de novo* structural-based comprehensive analysis of seven DDMCs and found their amino acid binding and coordination geometry propensities. We then utilized these results to characterize the correlation between metal selectivity, specific binding site composition and phylogenetic classification of the cation diffusion facilitator (CDF) protein family, a family of DDMC transporters found throughout evolution and sharing a conserved structure, yet with different members displaying distinct metal selectivity. Our analysis shows that DDMCs differ, at times significantly, in terms of their binding propensities, and that in each CDF clade, the metal selectivity-related binding site has a unique and conserved sequence signature. However, only limited correlation exists between the composition of the DDMC binding site in each clade and the metal selectivity shown by its proteins.

## Introduction

Divalent d-block metal cations (DDMCs) such as Zn^2+^, Mn^2+^, Fe^2 +^ and Cu^2+^, are essential for proper cell function. These metals play various roles in the cell, where they contribute to the stabilization of macromolecular structures and interactions and act as cofactors vital for enzyme activity. In many metal-dependent proteins, or metalloproteins, proper function depends on the presence of a specific metal cation. As such, the protein sequence, tertiary structure and the environment must contain elements that can differentiate between metal cations^1,2^. The specific chemical properties of each DDMC, such as size and electron configuration, dictate the composition and geometry of metal-binding sites and their immediate surroundings to afford selectivity and hence, better regulation of function^3^. DDMCs are usually bound to proteins via negatively charged residues (Asp, Glu) and polar residues (Cys, His and Asn, for example)^4^. Often, water molecules also participate in metal ligation and stabilize the metals in a particular conformation, while in other cases, different ligands bind or chelate the metals^5^. Although the residues that directly interact with cations (i.e., first shell residues) are crucial for metal binding, as well as for metal selectivity, it has been shown in several studies that second shell residues stabilize the first shell residues, dictate the specific polarity, size, geometry and dynamics of the binding site and are thus also required for appropriate binding of a metal^6,7^.

*Harding, Rulisek and Vondrásek, Zheng et al., Dokmanic´ et al*. and others scanned the RCSB Protein Data Bank (PDB) for metalloproteins prior to 2009 so as to map the binding properties of different metals, such as their tendency to bind specific residues and their coordination numbers^3,4,8–12^. *Laitaoja et al*. performed a specific analysis of zinc-binding properties based on PDB structures published before January, 2012^13^. As our research centers on understanding the metal-binding preferences of DDMCs to proteins belonging to the conserved family of cation diffusion facilitators (CDFs), we performed a *de novo* comprehensive scan of the RCSB PDB, filtering the relevant metalloproteins somewhat differently than was done in previous studies, with an emphasis on characterizing the binding properties of DDMCs.

The CDF family (TC# <u>2.A.4</u>) is a ubiquitous family, members of which are found in bacteria, archaea and eukaryotes^14^. In humans, CDF proteins are important for cell function, as revealed upon analysis of mutant CDF proteins (named Zinc Transporters (ZnTs)) that were found to be associated with severe diseases, such as Parkinson’s disease and type II diabetes^15,16^. Recent phylogenetic analysis divided CDF proteins into 18 clades, of which twelve were associated with a specific metal transport activity^17^. Each CDF protein specifically transports given DDMCs from the cytoplasm to the extracellular environment or into an inner cellular component, usually by exploiting the proton motive force and by undergoing conformational changes^18^. Accordingly, CDF proteins share a conserved structure, with the metal cations being transported through the transmembrane (TM) domain that contains the well-studied tight binding site, the A-site, composed of two residues from TM helix 2 and two residues from TM helix 5 (referred to as the quartet or XX-XX motif; Fig. 1A). The A-site is thought to have an important role in metal selectivity, as was shown in many studies where substitution of one of the quartet residues resulted in decreased protein function or a change in metal selectivity^18,19^. In *Escherichia coli* YiiP, for example, the only family member for which a crystal structure has been determined in an A-site-occupied state^20,21^, the A-site quartet is a DD-HD motif and is responsible for Cd and Zn transport. An exchange to HD-HD decreased Cd transport while maintaining Zn transport^22,23^. Although CDF proteins have been extensively studied over the past two decades, the A-site is the only metal binding site that was shown to be conserved within CDF proteins in terms of its impact on the function, while other sites, such as His-rich loops and the cytoplasmic domain, are not found in all CDF proteins^17–19^. Currently, the exact collective factors governing CDF proteins metal specificity remain elusive.

**Figure 1.**
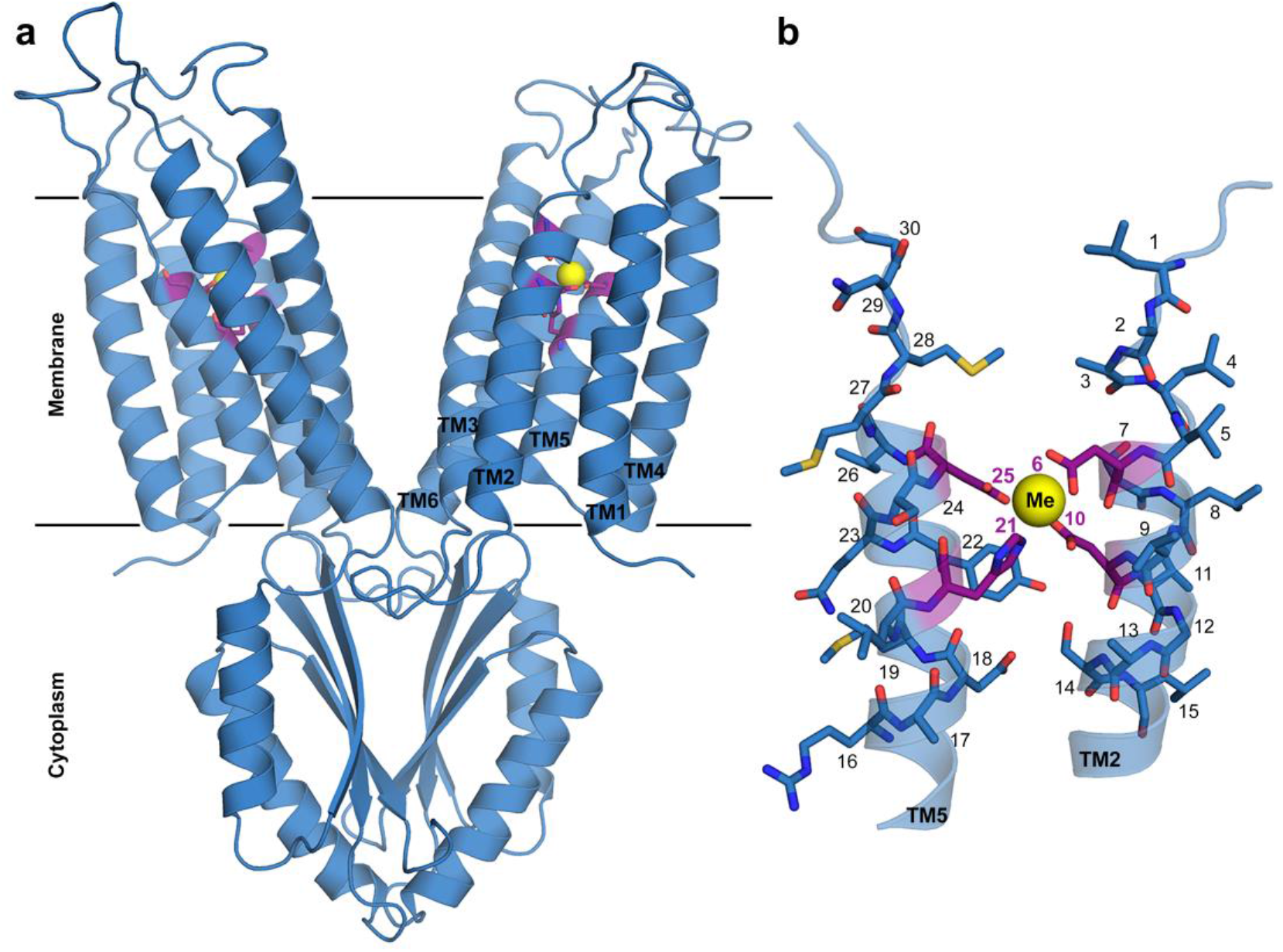
Structure of YiiP A-site. (**a**) Full structure of YiiP (PDB code: 3H90^20^) with A-site residues represented by purple sticks and Zn by a yellow sphere. (**b**) Magnification of TM2 and TM5 with A-site residues (purple) and their surrounding residues (cerulean blue) presented as sticks and numbered according to the legend to (the XX-XX quartet residues are numbered as 6, 10, 21 and 25 and the five residues up-and downstream to each X of the XX-XX quartet are numbered respectively). The metal cation is presented as a yellow sphere, nitrogen atoms are in blue, oxygen atoms are in red, sulfur atoms are in yellow and carbons are in purple or cerulean blue. Images were produced using PyMol^49^.

In this study, we aimed to understand if the A-site can globally control CDF proteins metal selectivity and if indeed there is a correlation between CDF proteins phylogenetic classification and metal selectivity, as previously proposed based on studies of some proteins in each clade^17,19^, studies that might not reflect the real specificity as usually not all DDMCs were tested and as *in vitro* studies might be misleading. More specifically, we sought to identify a unique signature of the A-site, including its immediate surrounding, in each CDF clade and to examine whether the clade A-site composition correlates with a specific protein function within the clade. To gain these insights, and as hundreds of proteins are assigned to the CDF family, we used comprehensive multidisciplinary computational approach. Accordingly, we combined our PDB-analysis with clade-specific multiple sequence alignment (MSA) and CDF proteins structural prediction to deeply characterize the relationships between CDF proteins phylogenetic classification, the A-site chemical environment and the experimental evidence for CDF protein metal preferences. We found that some metals have unique binding properties and that each CDF clade exhibits a distinctive A-site signature. However, in the case of CDF proteins, the correlation between A-site composition in each clade and its predicted metal selectivity holds only partially with the observed metal selectivity, probably due to structural features unique to this family.

## Results and Discussion

To investigate DDMC propensities, we analyzed metal-containing protein structures that had been deposited in the RCSB PDB up to June, 2016. Our analysis considered only those metals that were ligated by at least two protein-related atoms and allowed binding by only water molecules. Cofactors with specific coordination, such as heme molecules, were thus excluded from the selection. The inclusion of only water molecules as ligands is especially relevant for characterizing metal-binding in the CDF A-site, where the immediate surroundings are constrained by the tight intra membrane cavity to contain only water molecules, if any. Our analysis was based on X-ray structures with a resolution of 2.0 Å or better. These filtration criteria for metals in the database yielded the largest number for Zn, followed by Cd, Mn, Cu, Ni, Fe^3+^, Co and Fe^2+^ (Fig. 2, Supplementary Table S1 online; for full criteria, see *Methods*). This should be considered when referring to all of the results discussed, as it should be noted that for some metals, the pool is much bigger than others and, therefore, the binding probabilities for these metals are more confident. For iron, we considered Fe^2+^, as well as Fe^3+^, as Fe^2+^ tends to be oxidized in aqueous solutions. Indeed, even if Fe^2+^ was also added to the crystallization solution, there is no guarantee that this is the true iron oxidation state, unless experimentally confirmed. Fe^3+^ analysis thus serves as a reference for Fe^2+^. If the binding residues and geometries tendencies differ between the two iron forms, this then provides evidence for the true iron oxidation state.

**Figure 2.**
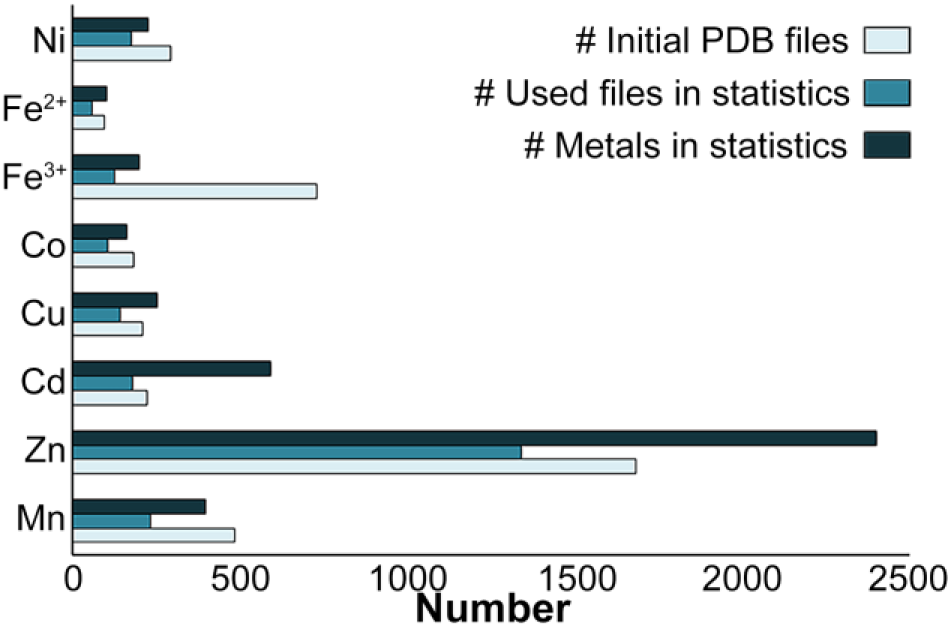
Number of used PDB files and metal cations in the analysis for each metal. The number of initial PDB files (light blue) refers to the number of files after primary filtration at the PDB website as described in Methods, while the numbers of used files (ocean blue) and metals (dark blue) considered in the statistical analysis refer to the final numbers after all filtration criteria were met.

The PDB-based comprehensive analysis of metal cations has some withdraws that need to be considered when analyzing the outcome results presented below: (1) The PDB website and files lack information on the method used to introduce the metal (soaking or co-crystallization with specific metals, or metal that was not added but was naturally bound to the protein in the cell), and if metal identity was confirmed experimentally. As the information can only be learned from the linked references, obtaining such data is problematic for metal meta-analyses. For example, in only 61% of 80 structures (10 for each metal) that we manually scanned, was the metal identity confirmed experimentally. Nonetheless, as DDMCs are less common in protein purification protocols, as DDMCs are not frequently found in cells and as DDMCs possess high electron density, even if not directly verified in the crystal form, one can be more confident of DDMC identification than of other ions such as sodium, potassium or chloride. In the case of meta-analysis, one can thus trust the database level-of-accuracy, even though this lack of knowledge lowers the confidence level of the results. (2) The metals bound in a crystal structure are not necessarily bound to the protein in the natural environment and do not necessarily reflect function. For example, metals might be bound due to their being part of the crystallization conditions or because use of a non-native metal resulted in successful crystallization, while use of the native metal did not (e.g., the crystal structure of a Mn-transporter that instead contains Zn). However, as proteins can be crystallized only when their structure is highly stable, metals bound to specific sites reflects such binding site as being favored thermodynamically. In the case of our analysis, metal-binding sites should be considered as they reflect an energetically favorable environment, although the data should not be over-interrupted in terms of function. (3) X-ray radiation can change the oxidation state of a metal during the crystal scanning. Major conformational changes in a crystal due to the different binding of different oxidation states of a metal are, however, less likely to produce data that can be processed. While structures with different oxidation states than stated due to radiation damage are less common, one should still take this error into consideration.

### Amino acid and coordination geometry analysis of different metals

The statistics of the amino acid, coordination number and coordination geometry propensities of all examined metals are summarized in Fig. 3 and Fig. 4. For amino acid and coordination number statistics, complete data are provided in Supplementary Table S2 online, while for complete coordination geometry data, see Supplementary Table S3 online. As expected from previous studies^3,4,8–12^, most metals are frequently bound by His, Asp and Glu residues and present an octahedral coordination geometry. Zn and Cu, however, present quite different preferences. While Zn has a high tendency to bind Cys with tetrahedral coordination, Cu binds almost exclusively to His, usually with trigonal coordination. Cd shows no statistically significant tendency for any of the coordination geometries but displays high preferences for the trigonal plane (with only two atoms bound), and for tetrahedral (not necessarily in full occupancy) and octahedral coordination geometries. Although the statistics show similar trends in the amino acid and coordination geometry analyses for some metals, each metal, nonetheless, presented a unique signature when both analyses were combined (Table 1).

**Figure 3.**
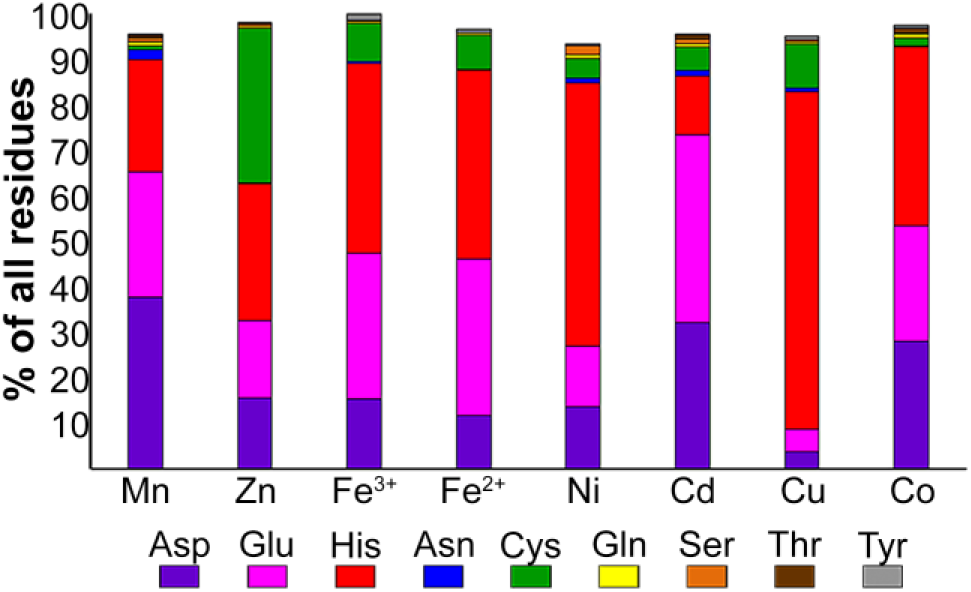
Selected amino acid propensities of the different metals. For each metal, the percentage of each residue is defined as the number of times that a specific residue was bound to the metal divided by the number of all residues bound to the metal. The different residues are presented in different colors: Aspartate in purple, glutamate in pink, histidine in red, asparagine in blue, cysteine in green, glutamine in yellow, serine in orange, threonine in brown and tyrosine in gray.

**Figure 4.**
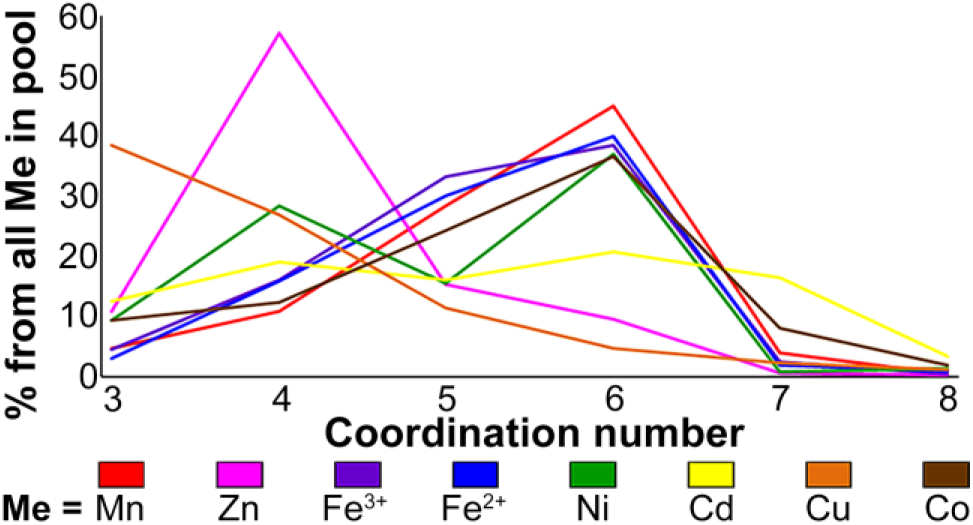
Main coordination number distribution of the different metals. For each metal, the percentage of each coordination number (CN, 3 to 8) is defined as the number of times the metal was bound in a specific CN divided by the total number of this metal in the statistics (in all coordination numbers). Every metal is presented in different color: Manganese in red, zinc in pink, ferric in purple, ferrous in blue, nickel in green, cadmium in yellow, copper in orange and cobalt in brown.

**Table 1.** The most abundant residues and coordination geometries (signature) of each metal.

| **Metal** | **Dominant residue(s)** | **Dominant geometry** |
| --- | --- | --- |
| **Zn** | Cys, His | Tetrahedral |
| **Cd** | Glu (Asp) | Diverse |
| **Mn** | Asp (Glu, His) | Octahedral |
| **Fe<sup>2+</sup>** | His, Glu | Octahedral |
| **Fe<sup>3+</sup>** | His (Glu) | Octahedral |
| **Co** | His (Asp, Glu) | Octahedral |
| **Ni** | His | Octahedral |
| **Cu** | His | Trigonal plane |

In our approach, we counted the amino acids for each atom that ligates the metal (if an Asp ligates the metal with both side chain oxygen atoms, this Asp was counted twice). To verify that this method is not biased towards residues with more than one atom ligating the metal (e.g. Asp, Asn, etc.), we also calculated propensities by residues rather than by atoms. As can be seen in Supplementary Fig. S1 and Table S4 online, this approach yielded similar trends, with only minor numerical changes.

Further to these analyses, we wanted to establish a subset of membrane proteins that will allow more accurate reference when considering the results for the CDF specific study. However, as to May, 2017, our search for membrane proteins in the RCSB PDB revealed that only 18 DMTC bound in TM regions are found (when filtering symmetrical or homologous binding sites and using a resolution cut-off of 2.5 Å to increase the pool of metals but to use data that is still reliable). With 11 metal-binding sites for Zn and 0-2 sites for other metals, this analysis cannot be performed efficiently.

Our results are in partial agreement with previous analyses showing that binding residue distribution for each metal is sometimes different (for example, *Dokmanic´ et al*.^3^ reported that Ni binds more Cys than Asp and Cd more Cys than His, a trend opposite to what we find in our analysis), although in most cases, the distribution was similar, which also holds true for coordination number distribution. However, since the metal filtering criteria used were different in each study, the results cannot be compared directly. For example, Rulisek and Vondrásek^4^ did not limit resolution, used metals that are bound by ligands (unless DNA), differently defined the atoms that are bound to each metal (by distance rather than by LINK record), did not state whether a non-redundant set of proteins was used and considered a very small pool of metals used (maximum 49). *Zheng et al*.^12^ used a cut-off resolution of 1.5 Å or better and a non-redundant data set for analysis of residue binding, and did not limit the resolution and redundancy for coordination number calculations. Finally, *Dokmanic´ et al*.^3^ used a cut-off resolution of 1.5 Å or better and a non-redundant set of proteins but also used metals that bind ligands and defined the bound atoms (only S, N, O and Cl) in terms of distance from the metal. Probably the most influential criteria that led to somewhat different patterns than reported in previous studies were the restrictions imposed on the number and type of metal-bound ligands. Nonetheless, our results, like those of previous studies, did not, for the most part, reveal significant differences between the different metals. Although, in some cases, the coordination number or residue distribution were unique to a metal (for example, the specific very high tendency of Cu to bind His and the tetrahedral coordination that characterizes Zn-binding), generally speaking, such tendencies appeared to be quite similar in different metals, with most metals being coordinated in an octahedral geometry by Cys, Asp, Glu and His, with no clearly significant preference for any of these residues. Since even small differences in tendencies may relate to affinity, this analysis can help in the design of metal-binding sites with a higher probability of binding a specific metal. While such analysis is not sufficient for predicting metal selectivity in metalloproteins, it can help explain (or contradict) known or putative trends.

### pH-dependence of metal propensities

The protonated state of an amino acid changes in response to changes in the surrounding pH, with the most physiologically relevant change being that of His, with a pKa of ~6.04^24^. This is particularly important when considering CDF proteins, as most are thought to rely on a mechanism involving proton exchange, such as shown to take place over the A-site motif histidine in YiiP^25^. A pH-based analysis can contribute in individual CDFs’ investigations or modifications, and can give insights to variety of studies of metalloproteins binding abilities. Yet, as the phylogenetic classification is not related with the pH, this analysis is less applicable in this study. However, as such analysis was not considered in previous similar studies, we decided to utilize our data to examine whether a pH-dependent bias in metal ligation and coordination exists. For that, we divided the analyzed metals based on their pH, namely lower or equal to 6, between 6 and 7, between 7 and 8 or 8 or higher. The data are summarized in Supplementary Fig. S2 and S3 online (complete data are presented in Tables S2 and S3). As expected, in all metals, His is more dominant in structures with basic pH values, as compared to its average tendency, consistent with lower tendencies of Asp and/or Glu to appear in such roles. Nevertheless, the PDB-based pH-dependent amino acid, coordination number and coordination geometry preferences should be analyzed with caution, mainly because the pH listed in PDB files is not accurate and usually refers to the crystallization solution rather than the overall pH in the crystallization vial (which also includes the protein and its storage solution, at the least) nor necessarily reflects the pH during actual metal binding.

### Compositional analysis of metal quartets

Since the CDFs A-site is composed of four residues that participate in metal binding, we analyzed the distribution of different quartets of amino acids for each DDMC in the PDB. For all metals addressed in such analysis, we considered only those that were bound to exactly four residues (and as such, bound to at least four atoms), with or without water molecules. Fig. 5 and Supplementary Table S5 online show clear quartet preferences for some metals when bound to four residues. For example, Fe^2+^ shows higher tendency to be bound by EEEH and lower to DHHH, as compared to Fe^3+^. Excluding cysteine-containing quartets, which are more likely to be involved in stabilizing structures rather than directly contribute to enzyme activity (and, therefore, together with the fact the cysteine residues are less found in CDF quartets, are less relevant for our CDF analysis)^13^, Zn is frequently bound to DHHH, Cu to HHHH and Cd to negatively-charged quartets, like EEEH. Mn tends to be bound by DHHH as well and Co and Ni are both highly bound to the EHHH quartet, while Ni tends to bind more His-containing quartets, such as DHHH and DDHH, and Co is usually bound to more charged quartets, with high tendency to be bound to DEEH and DDEH.

**Figure 5.**
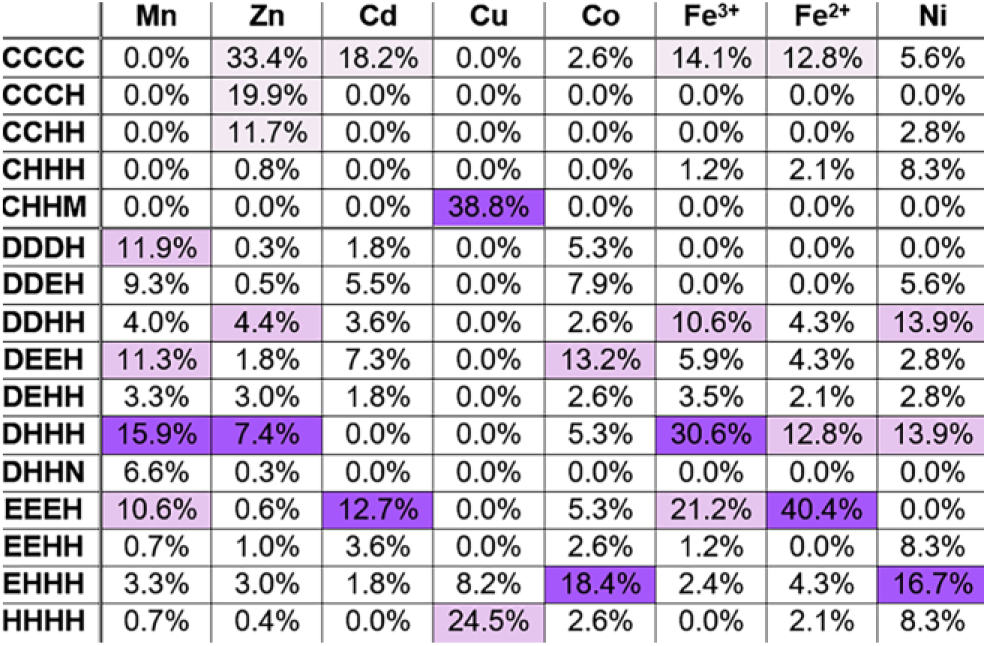
Most abundant quartet distribution of the different metals. For each metal, the quartet percentage is defined as the number of times the metal was bound by the specific quartet divided by the total number of quartets to which the metal was bound. The cysteine-containing quartets (first five lines) are separated from the other quartets due to the tendency of the former to participate in structural sites rather than enzyme active sites. Darker purple colors represent higher binding tendencies. As they are less relevant for enzymes active sites, the high tendencies of quartets containing more than one cysteine (first three lines, not including Cu) are exceptional and are presented in light purple.

### Analysis of the A-site of CDF proteins

In our study, we examined the relationship between metal specificity and the phylogenetic classification of CDF proteins proposed by *Cubillas et al*.^17^. Specifically, we sought to understand, based on our PDB analysis, whether the composition of the CDF A-site in each clade could determine the metal selectivity of the proteins in that clade. To do so, we characterized A-site composition in each clade in search of unique clade-related signatures. This was achieved by clade-specific MSA against the YiiP sequence to identify the A-site residues (Supplementary Table S6 online). Such information was used to generate LOGO presentations of binding site residues and their immediate neighbors (i.e., five residues from the N- and C-termini of each residue of the XX-XX quartet). Recognition of the A-site was confirmed by structural modeling of a representative protein from each clade and overlapping that structure on the solved YiiP structure (PDB # 3H90, bound to Zn^20^). The LOGO sequences (Fig. 6) show clear differences between the clades. The most dominant quartet is HD-HD, found in eight clades. Since the PDB-based quartet analysis showed the DDHH quartet to be common for all metals except Cu, it is not surprising that this quartet was also common to A-sites in CDFs. Unique quartets were ND-DD in clade 1, DD-DD in clade 4, DD-HD in clade 5, EN-HD in clade 6 and HH-DH in clade 12. Although the HD-HD quartet showed up in many clades, each clade presenting this quartet also contained a unique signature of second-shell residues that can influence binding site properties such as polarity, and hence could play a role in clade-specific metal selectivity. For example, clade 2 contains Asp at position 14, Asn at position 18 and His at position 22, while clade 3 contains Asn at position 6 and Thr at position 13 (positions defined in Fig. 1B).

**Figure 6.**
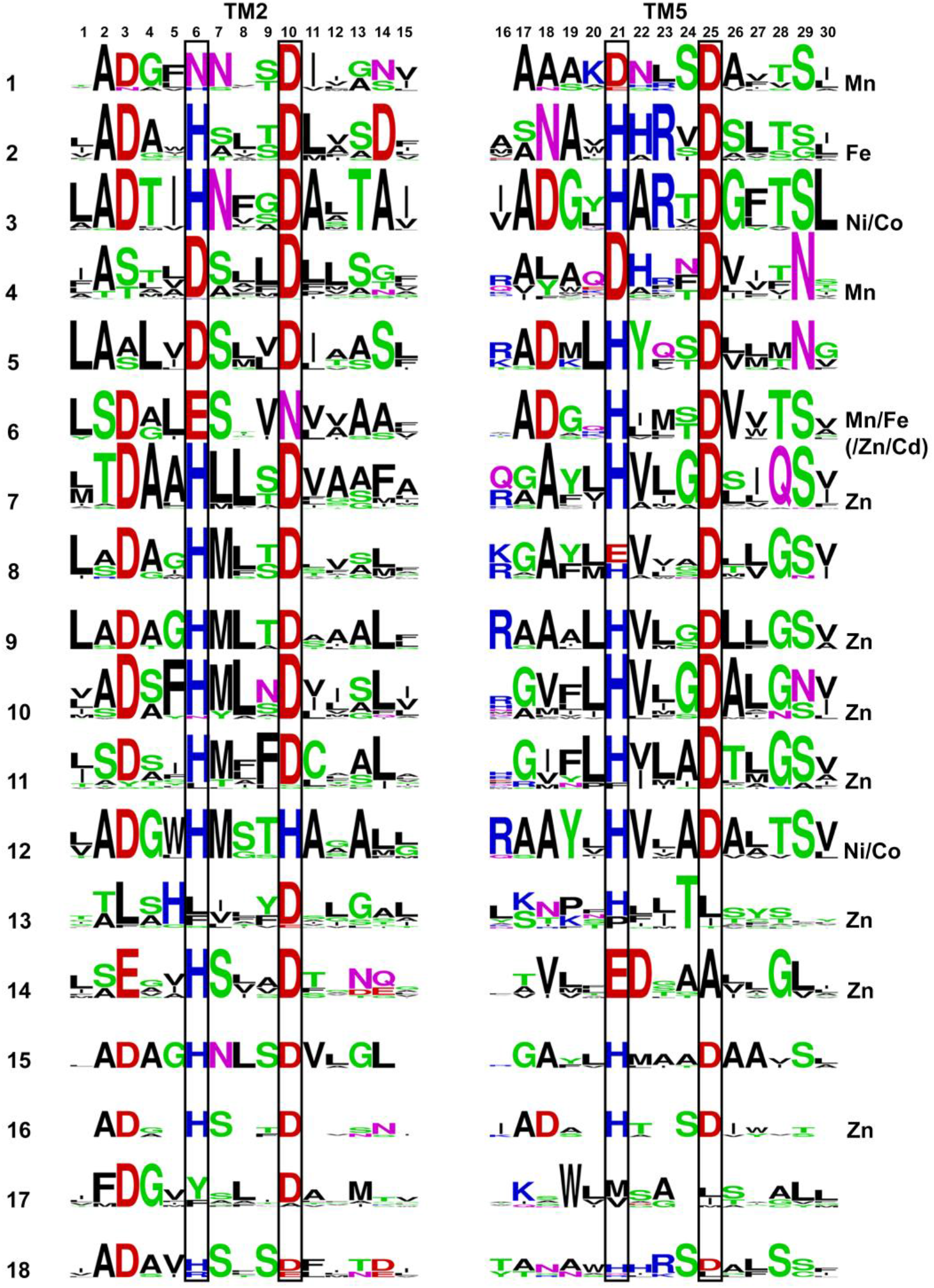
LOGO presentations of all CDF clade A-sites (all at the same scale with a maximal height of 4 bits). Each 15-residue block contains the A-site residues in the TM helix in black boxes plus five residues from the N- and C-terminus. Position 6 overlaps with D45, position 10 with D49, position 21 with H153 and position 25 with D157 in YiiP. Acidic residues (E, D) are presented in red, basic residues (H, K, R) are presented in blue, small/polar (G, C, S, T, Y) are presented in green, amide residues (N, Q) are presented in pink and hydrophobic residues (A, L,V, I, M, P, F, W) are presented in black. Metals shown experimentally to be preferably transported by proteins in a clade are indicated for each clade on the right.

### Correlation between phylogenetic classification and A-site conservation

We next wanted to understand whether the A-site specifically is more conserved within different domains of life, or between clades which are evolutionary close, compared to more distant clades. Clades 4, 7, 10, 11 and 13 contain only eukaryotic proteins, clades 1, 5, 6, 8, 9, 12, 15 and 17 contain only bacterial proteins, clade 18 contains only archaeal proteins and clades 2, 3, 14 and 16 include mixtures of eukaryotic, archaeal and/or bacterial proteins. When we analyzed the bacterial and eukaryotic domains separately, we saw that there is no fully conserved position found only in a specific domain, relative to the others. Specifically, A-site residues vary between the clades in each domain, and in the bacterial-related domains, other than clades 9 and 15 that both contain the same HD-HD quartet, each domain contains a different quartet (Fig 6 and Supplementary Table S6 online). This trend might hint at a relationship between phylogenetic classification and the related quartet composition to different metal selectivity, in each domain.

The closest clades in terms of evolutionary distance are the bacterial clades 8 and 9, both of which were assigned as Zn-clades although clade 8 was also shown to transport other metals^17^. The clade 9 quartet is consistently HD-HD, while the proteins in clade 8 contain either HD-HD or HD-ED quartets, yet proteins that are more distant from clade 9 contain the HD-ED motif. Other than this position and position 19 (Fig. 1B and 6), all the other positions exhibit similar compositions. Clades 7-11 are uniquely grouped together with all clades at very short distances^17^, all contain the HD-HD motif (except of clade 8 with HD-(E/H)D), show similar general composition and very similar hydrophobic signatures between each quartet couple (positions 7, 8, 22 and 23, Fig. 6). All the other quartets are more distant from each other and show unique signatures in the A-site, if not in the quartet residues than in the second shell residues. However, in some cases, there are as many differences between close clades as for between distant clades, meaning that the distances are only partially reflected by the A-site. If a correlation between A-site composition of each clade and metal selectivity indeed exists, evolutionary pressure may have caused more drastic changes in the A-site in some close clades, so as to enable versatile selectivity.

### Correlation between selected CDF clade A-site signatures and metal selectivity

Montanini *et al*. performed in 2007 similar attempts to ours, to find the relationship between the phylogenetic classification, A-site composition and metal selectivity, and suggested that the A-site might have a role in metal selectivity and that this site should be a starting point for further investigation^19^. Additionally, Cubillas *et al*.^17^ suggested a correlation between phylogenetic classification and metal selectivity. However, their work contains cases where only few metals were associated to the same clade, or very little experimental evidence for specific metal-transport in each clade is available (see Supplementary Table S6 online). Although Montanini *et al*. used similar approaches in their analyses^19^, our analyses are different and updated than theirs, considering the data available today: we used structural data that was not available at the time, different and deeper phylogenetic classification that contains larger number of analyzed subfamilies, examination of larger area of the A-site (including 26 second-shell residues, compared to 6), use the metals structural-based, CDF-specifically-designed comprehensive-analysis and more experimental data from recent years. Overall, although the PDB-based metal analysis showed different patterns in metal preference and although each clade shows a unique A-site composition, we observed that there is no good correlation between the A-site composition in each CDF clade, the composition ability to prefer specific metal binding and the experimental evidence for the clades’ metal specificity. We discuss below selected clades and show how, in some cases, the general metal analysis can well explain metal selectivity observed experimentally, while in other cases, it does not.

Clades that were experimentally shown to *mainly* transport Zn (more than one protein per clade; the evolutionarily-closed clades 7, 10 and 11^17^, Supplementary Table S6 online) contained the HD-HD motif, in which the first His was previously shown to control Zn selectivity over Cd in the ZnT and YiiP proteins^22^. As Zn requires a lower coordination number than do other metals, A-sites with multiple His residues, which allows pH-based regulation and proton transfer, are sufficient for Zn stabilization in the A-site. Our metal analysis results show that in contrast to Zn, Cd had the lowest probability of all the transition metals to bind His and high probability to bind Glu and Asp (Fig. 3). Indeed, YiiP (clade 5), that transports both Zn and Cd, the latter with higher affinity^23^, contains the DD-HD motif (and hence allows also bigger cavity space), which might reflect a compromise between both ions with a bias towards the bigger Cd.

Proteins in clades 1 (bacteria) and 4 (eukaryotes) that share a quite recent common ancestor^17^, were experimentally shown to display selectivity to Mn^17^ (Supplementary Table S6 online). Although their A-site compositions differ (corresponding to ND-DD and DD-DD, respectively), clade 1 contains Asp and clade 4 contains a conserved Asn at putative loops at the end of the TM helices that face the side opposite the cytoplasm (Fig. 1B and 6) and they, hence, might play a role in the uptake of the incoming ion. All proteins that were experimentally shown to be involved in manganese homeostasis outside of these clades also contain Asn in the A-site motif^26– 29^. This is in agreement with our metal analysis showing that Mn tends to bind Asp and Asn more than other metals (Fig. 3) and in a coordination geometry that is possible with multiple Asn/Asp residues (octahedral). Taken together, these results suggest that A-site quartets composed mainly of Asn and Asp, and with Asn and Asp in the immediate surroundings, were selected through evolution to participate in manganese stabilization.

Clade 2 was shown to be mostly related with iron transport^17^ (Supplementary Table S6 online), and although its A-site composition is the same as for Zn clades (where the HD-HD motif is found), this clade also contains a conserved and unique Asp at position 14 and an Asn at position 18, both of which are found in the inner-TM side of the A-site (Fig. 1 and 6). Since both Zn and Fe^2+^ tend to bind His, the HD-HD motif can trap better these metals than other metals, while Asp14 and Asn18 might help to coordinate Fe^2+^ in a preferred geometry. Since Fe^2+^ has a low preference for binding Asp, our results offer only limited explanation for the composition of this A-site. We thus suggest that in this case, the composition of the A-site region is not solely responsible for metal selectivity.

Clade 6 (EN-HD motif) was shown to be mainly related with Mn and Fe^2+^ transport but also with Zn and Cd^17^ (Supplementary Table S6 online). As Fe^2+^ and Cd have greater probability to bind Glu, Zn to His and Mn to Asp, A-site composition might be important for attracting all of these ions, serving as a non-specific metal-binding site. Interestingly, in our PDB-based quartet analysis, no metal was bound to the DEHN motif.

### Clade 4 as an example of an incoming-ion effect on A-site composition

As described above, the A-site of CDF proteins contains four residues that are assumed to take part in metal binding. Considering that CDF proteins are antiporters, it is highly important when examining A-site composition to also consider the incoming ion. Although most CDFs are thought to exploit the proton motive force to enable transition metal cation transport, it has been shown that the potassium gradient can also be utilized for this purpose^18,30,31^. Since the TMD is highly hydrophobic, it is only reasonable to assume that the incoming ion is transported through the same hydrophilic conserved site and, therefore, its composition is possibly also related to the identity of the incoming ion. For example, in *E. coli* YiiP, which uses the proton motive force for metal transport, the proton moves through the His of the DD-HD motif^25^. Although it was shown previously that acidic residues can participate in proton movement that is coupled to cation transport (for example, in Na^+^/K^+^-ATPase^32^), in the case of CDF proteins with A-site quartets that do not contain His, one should also consider other transport mechanisms or incoming ions.

DD-DD A-site composition in clade 4 reflects a unique, highly charged site. Our DDMC PDB-based analysis showed that no metal tends to bind to a site with this composition. Therefore, we performed specific analysis of the DDDD quartet for all of the metals considered, using the same criteria but with no resolution or method limitations, in order to find evidence for metal preference. This analysis did not uncover any new structures, as compared to our previous analysis, meaning that there is only one Zn that is bound by the DDDD quartet in the PDB. Experimentally, only manganese homeostasis has been shown to be related to proteins in this clade (AtMTP8^33^, ShMTP1 and AtMTP11^17^) and, indeed, our analysis showed that Mn has the highest propensity for Asp-binding. Although to date only protons and potassium have been experimentally shown to serve as incoming ions, other ions should be considered as well, as CDF proteins are found in both cell and intracellular compartment membranes. Regarding the His-lacking highly charged DD-DD motif, one should address not only mono- but also divalent cations of considerable sizes, such as Ca^2+^ and Mg^2+^, as better stabilizing this site. Since K^+^ and Na^+^ are abundant in protein purification and crystallization protocols and given how they present lower charge and less defined coordination than do divalent cations, binding of K^+^ and Na^+^ to proteins in crystal form can be non-specific and misleading. Therefore, we performed the same limit-free resolution analysis as conducted for transition metal cations for Mg^2+^ and Ca^2+^ to examine whether they tend to be bound at DDDD sites. Such analysis compared similar numbers of structures for both cations, yet found that only Ca^2+^ ions were bound to this site. With 24 hits, this is the most abundant metal bound to the DDDD quartet. Since Mg^2+^ ions have greater electronegativity and thus a larger hydration shell, they are less likely to be transported through the intramembrane site in CDFs (for example, Mg^2+^ has been shown to block Ca^2+^ channels^34^). Ca^2+^, with the same charge and similar size to transition metals, and with an intracellular concentration roughly 10,000 times lower than its extracellular concentration^35^, is more likely to be the force that drives transition metal transport from the cytoplasm to the extracellular environment. In support of this hypothesis, it was recently shown that a DDDD site is related to Ca-selectivity in the TRPV6 Ca^2+^ channel^36^. All of this suggests that in the case of the eukaryote clade 4 and the DD-DD motif, one should consider that specificity might be related to an incoming Ca^2+^ ion and not only to the transition metal cation, but this should be examined *in vivo* and *in vitro*. In light of these results, we speculate that A-site composition also determines incoming ion identity.

### Concluding Remarks

Our analyses demonstrate differences between metal cation propensities, although as the number of residues and geometries are limited, the observed degree of versatility is low. Our results further show that the composition of the immediate environment of the A-site can be related to metal selectivity only at the single-protein level and suggest that when considering this site, the phylogenetic classification of CDF proteins is not related to their metal selectivity. Numerous studies have shown that even a single amino acid substitution in the A-site XX-XX motif changed the metal selectivity of a CDF protein (for example, MntE^37^, ZnT proteins^22^ and YiiP^23,38^), thus emphasizing the importance of this motif for metal specificity. Yet, both previous experimental findings and our current results raise several points that need to be taken into consideration in CDF protein phylogenetically-related metal selectivity studies.

First, different clades with the same quartet motif show different metal selectivity. As such, other elements in the protein, such as second shell residues, the flexibility of the TMD that can influence the geometry of first shell residues, or other metal-binding sites, must also be related to the specificity seen. Our results propose that in some cases the second-shell residues may have an impact on metal selectivity (clades 1 and 4 for example), but all these factors should be examined at the single-protein level. Second, while in some clades all of the proteins that were tested displayed the same specificity, in other clades, different proteins presented different metal selectivity. Moreover, only in few clades was there significant evidence for metal specificity, pointing to a lack of any strong indication of the genetic relatedness of metal specificity in CDF proteins. Hence, based on the current knowledge in the field, we hypothesize that only proteins that are affiliated with specific clades might share the characteristic selectivity of that clade (specifically, proteins in clade 1, 2, 4, 7, 8, 11 and 12). Third, in the majority of the CDF protein studies, not all DDMCs were tested, meaning that experimentally-based metal selectivity classification might be misleading. Furthermore, *in vitro* analysis might not reflect the true biological role of the proteins studied. Fourth, many CDF proteins have been experimentally shown to transport different metals, so that if each metal binding has a different Kd, as was demonstrated for YiiP^23^, this would mean that CDF proteins can play diverse roles and possess different selectivities, depending on the environmental conditions. Lastly, as speculated here, CDF metal selectivity might, in some cases, be related to the nature of the incoming ion and not to a specific transition metal cation. Accordingly, one should consider the incoming ion identity when studying the mechanisms of CDF proteins.

## Methods

### Metals selection for statistics

The experimental data, including the coordination file, of every atomic structure that was determined and published, is deposited in the RCSB PDB (http://www.rcsb.org/pdb/home/home.do). For each metal (that was shown to be transported by CDF proteins, aka Cd^2+^, Co^2+^, Cu^2+^, Fe^2+^, Mn^2+^, Ni^2+^ and Zn^2+^, as well as Fe^3+^), the following criteria were used for filtration of PDB coordination files listed as of the end of May, 2016: (1) The deposited structure had to contain protein (and not only DNA for example), (2) X-ray structures had to be of a resolution of 2.0 Å or better (so no electron microscopy or NMR structures were used, and to use only structures with good confident in their metal assignment), (3) the PDB file had to contain a LINK record with the atom label (for MN, ZN, NI, FE (Fe^3+^), CO, CU, CD) or component (for Fe^2+^, FE2) in a metal coordination connection, meaning a metal is coordinated by organic molecule in the structure. For structures with more than 90% sequence identity, only a representative structure was retrieved, so if for the same protein there are different structures (mutants, different space groups, etc.), only one of them was considered in order to avoid biased results.

Subsequently, for each of the extracted structures, if there were symmetry-related metal-binding sites, only one representative binding site for each metal was considered. If any ligand other than water (such as other metal ions, DNA molecules or other small organic ligands) was bound to a metal cation, this metal was omitted from the statistics. To decrease bias due to non-specific binding, only metals bound by at least two protein-related atoms were considered. The number of extracted metals in each filtration step can be found in Supplementary Table S1 online.

### Propensity analysis of amino acids and coordination geometry

For each metal that was considered in our statistical analysis, we checked which atoms from each residues were ligated and what was the coordination number. In the amino acid preferences analysis, we counted those atoms that are involved in metal coordination rather than residues (if Asn chelated the metal with both O and N, Asn was counted twice), unless otherwise specified. The metal-bound atom identification relied on the relevant LINK record in the PDB file (the file contains an entry for each linked metal-atom pair). For each metal, the percentage of a specific residue was calculated as the number of times the metal was found bound to this residue (total number of bonds to the residue’s atoms from all used metals), compared to the total number of atoms that were bound to all the specific metal cations (total number of bonds to all of the residues’ atoms from all used metals). Coordination geometries for all metals considered were calculated by the stand-alone version of the FindGeo server in its default settings^39^. FindGeo accepts the metal and coordination ligands (a PDB file) as an input. The output is the most probable geometry for metal coordination from a library that includes a total of 36 ideal coordination geometries (coordination number 2-9). Metals with coordination geometries that do not correlate with any of the library geometries were not considered. For each metal, the percentage of a specific coordination number/geometry was calculated as the number of times the metal was found bound at this coordination number/geometry, compared to the total number of metals in the pool (all metals in all coordination numbers or in the pool of metals with defined geometry, respectively).

The pH values for the pH-dependent analysis were extracted from the experimental procedure recorded in the PDB file (under REMARK 200). Files that were missing this information were considered only in non pH analyses.

### Multiple sequence alignment and LOGO sequence

CDF proteins assignments into clades and their sequences were extracted from *Cubillas et al*.^17^. For each clade, MSA was performed on all proteins assigned to that clade and the *E. coli* YiiP sequence, the only CDF proteins for which atomic resolution structure in the A-site bound state exists. MSA was performed using ClustalO^40,41^, while LOGO presentations of each A-site residue, as well as the five residues up- and downstream of each X of the XX-XX quartet, were generated by WebLogo^42^. For clades 13, 14, 17 and 18, the clade-specific MSA failed to predict the second duet residues comprising the A-site to be of a composition suitable for metal binding. Therefore, MSA with CDF protein sequences from all clades was performed and the A-site composition was extracted.

### Structural models

Structural modeling of one representative protein from each clade was performed using the SWISS-MODEL Automatic Modelling Mode^43–46^ and MPI Modeller^47^ based on the YiiP structure in the Zn-bound state (PDB # 3H90^20^). The representative sequence for modeling each clade was selected using the pairwise identity score matrix calculated by ClustalO. For each protein, both models were overlapped and their quality was manually assessed; best model (no clashes, more stabilizing interactions, etc.) was chosen for further investigation. Verification of the MSA-based A-site residues was performed by overlapping the models on the YiiP structure using the iterative magic fit and fragment alternate fit tools in Swiss-PdbViewer 4.1.0^48^. For clade 17, as the MSA-based A-site and the structure-based A-site did not overlap, the MSA-based A-site composition is listed here as the structure-based site was not conserved within the clade.

### DDDD quartet analysis

For each metal (Cd^2+^, Co^2+^, Cu^2+^, Fe^2+^, Fe^3+^, Mn^2+^, Ni^2+^, Zn^2+^, Ca^2+^ and Mg^2+^), the same parameters were used as in the metals selection for statistics method, above, except that (1) filtration was conducted at the end of June, 2016 for all metals, (2) the LINK record had to also contain a connected component ASP (to verify that only structures with metals that are bound to at least one Asp residue will be considered), and (3) the analysis did not include resolution or experimental method limitations (in order to increase the pool). Such analysis yielded the following numbers of PDB files: Cd^2+^, 250; Co^2+^, 131; Cu^2+^, 43; Fe^2+^, 82; Fe^3+^, 271; Mn^2+^, 844; Ni^2+^, 153; Zn^2+^, 1278; Ca^2+^, 2310; Mg^2+^, 2538.

### Data availability

The datasets generated during and/or analyzed during the current study are available from the corresponding author on reasonable request.

## Acknowledgments

We thank Dr. Ciro Alberto Cubillas for providing CDF protein sequences and Dr. Claudia Andreini for her help with FindGeo program. The authors of this work are supported by the Israel Ministry of Science, Technology and Space, the Israel Science Foundation (grant no. 167/16), the European Molecular Biology Organization and CMST COST Action CM1306.

## Author Contributions

All authors designed the study and wrote the paper. S.B.Z. extracted the data and performed the meta-analyses. All authors analyzed the results and approved the final version of the manuscript.

## Additional information

### Supplementary information

Two electronic supplementary information files accompany this paper: ESI1 contains supporting information figures, ESI2 contains supporting information tables.

